# PRLX-93936 and BMS-214662 are cytotoxic molecular glues that leverage TRIM21 to degrade nucleoporins

**DOI:** 10.1101/2024.12.18.629219

**Authors:** Marc A. Scemama de Gialluly, Anthony R. Allen, Elijah H. Hayes, Patrick Zhuang, Ralston B. Goldfarb, Amanda N. Farrar, Yuriy Fedorov, Drew J. Adams

## Abstract

Although molecular glues have emerged as innovative tools within the field of chemically induced proximity, approaches for their discovery remain limited. Here we report a phenotypic screening approach in which small molecules whose cytotoxic mechanisms require ubiquitination show gain of viability following pharmacological inhibition of the E1 enzyme UAE. This approach revealed two previously clinically-evaluated cytotoxins of unknown mechanism, PRLX-93936 and BMS-214662, as molecular glues that directly target the E3 ubiquitin ligase TRIM21. These molecules induce TRIM21-mediated proteasomal degradation of multiple nucleoporin proteins, leading to inhibition of nuclear export and ultimately cell death. Loss of nucleoporins and nuclear export accounts for past observations in which BMS-214662 led to disrupted subcellular protein localization. Furthermore, the cytotoxicity of these agents correlates strongly with TRIM21 expression, suggesting clinical re-evaluation of these agents in patients with TRIM21-high cancers. Additionally, relative to recently-reported TRIM21-targeting glues, these two scaffolds display high cellular potency, creating new opportunities for targeted protein degradation via the design of additional glues and ‘TRIMTACs’.

## Introduction

The mechanistic elucidation of small molecules like lenalidomide, indisulam, and CR-8 as ‘molecular glues’ that induce an E3 ligase to ubiquitinate and degrade a target protein has led to a surge of interest in targeted protein degradation (TPD).^1–7^ This concept has led to identification of additional molecular glues and proteolysis targeting chimeras (PROTACs), which comprise an E3-targeting small molecule, a linker, and a target protein ligand to achieve TPD.^8,9^ While molecular glues remain challenging to discover, recent work has begun to define design strategies and screening approaches capable of identifying molecular glues in a more systematic way.^10–14^ Recently we reported a variant of an established approach^15^ in which pharmacological inhibition of Cullin RING ligases (CRLs) could reveal small molecules whose cytotoxic mechanisms involved CRL-mediated protein degradation.^16^ While CRL family substrate receptors like cereblon, VHL, and DCAF15 mediate most molecular glues and PROTACs identified to date, E3 ligases represent a large and biologically diverse protein family, and expanding the repertoire of E3 ligases that can be leveraged for TPD remains of broad interest.^17,18^

Here we implemented a screening strategy in which pharmacological inhibition of all E3 ligase complexes revealed PRLX-93936, a cytotoxin of unknown mechanism of action,^19,20^ as a novel molecular glue targeting TRIM21. TRIM21, a member of the TRIM family of E3 ligases, has been well studied in immunology, where it is known to oligomerize and ubiquitinate immunoglobulin G (IgG)-coated pathogens to mediate their rapid degradation.^21–24^ The ability of TRIM21 to degrade antibody-bound cargoes has also been harnessed for the TRIM-AWAY platform, in which cellular delivery of an antibody against a target protein can induce its TRIM21-mediated degradation.^25,26^ Recent reports have also explored the therapeutic potential of this approach by delivering viruses encoding TRIM21-nanobody fusion proteins targeting aggregation-prone proteins including Tau.^27,28^ More recently, the first molecular glue targeting TRIM21, (*S*)-hydroxy-acepromazine, was reported and shown to induce degradation of nuclear pore proteins and impair nuclear transport.^29^ An additional series of TRIM21-targeting glues was also recently reported.^30^ Excitingly, hydroxy-acepromazine also was elaborated to PROTACs (or, ‘TRIMTACs’) that selectively induced degradation of target proteins when they were forced into protein aggregates or condensates but not when the target protein was soluble. This finding likely reflects TRIM21’s biological preference for ubiquitinating large multimeric complexes and suggests TRIM21 could be an optimal E3 ligase in contexts where clearance of protein aggregates could be therapeutic.^29^

Here we identify PRLX-93936 and BMS-214662 as two novel series of TRIM21-targeting molecular glues. These molecules are structurally unrelated to hydroxy-acepromazine and are orders of magnitude more potent when tested in head-to-head assays. Like hydroxy-acepromazine, PRLX-93936 and BMS-214662 induce rapid degradation of a wide swath of nuclear pore proteins, leading to loss of nuclear trafficking and cell death. We also confirm that genetic disruption of NUP98’s autoproteolysis domain effectively prevents PRLX-93936-mediated nucleoporin degradation and cell death, further supporting a direct glue mechanism between TRIM21 and NUP98. Together these studies define the anticancer mechanism of PRLX-93936 and BMS-214662 and provide high-quality chemical matter to propel the development of novel TRIM21-targeting small molecules for targeted protein degradation.

## Results

Recently we reported a screening approach that used pharmacological inhibition of neddylation to identify small molecules whose mechanism of cytotoxicity requires Cullin RING Ligase activity.^16^ A limitation of this approach is that only the Cullin RING subfamily of E3 ligases are inactivated by inhibiting neddylation. We imagined that inactivating a wider range of E3 ligases could potentially spotlight additional degraders whose mechanism targets non-CRL E3 ligases. Small-molecule inhibition of the ubiquitin activating enzyme (UBA1/UAE) prevents activation of a large majority of E3 ligases and is currently being pursued as an anticancer strategy.^31,32^ While sustained treatment with the UAE inhibitor TAK-243 was potently cytotoxic to the acute myeloid leukemia cell line OCI-AML-3 (EC_50_ 15 nM), treatment for 10 h was better-tolerated. Short-term treatment with TAK-243 thus provided a meaningful window during which cells could be maintained in a viable state without E3 ligase activity (Supp. Fig. 1a).

We next established a comparative cell viability assay in 384-well plates in which the impact of small molecules was compared with and without addition of TAK-243; we then evaluated 2,000 known bioactive small molecules using this assay (Fig. 1a,b). Only 5 hits were identified that showed <50% viability and also gained at least 30% viability with TAK-243 co-treatment (Fig. 1b). Hit validation across a wide concentration-response range confirmed only one of the five, PRLX-93936, as having diminished cytotoxicity when co-treated with TAK-243 (Fig. 1c,d; Supp. Fig. 1). This low confirmed hit rate likely reflects both the uncommon nature of ubiquitination-dependent cytotoxic mechanisms and the requirement for induction of cell death within 10 hours. PRLX-93936 cytotoxicity could also be rescued by the proteasome inhibitor bortezomib, supporting a mechanism of cytotoxicity dependent on the proteasome (Fig. 1d).

**Figure 1.**
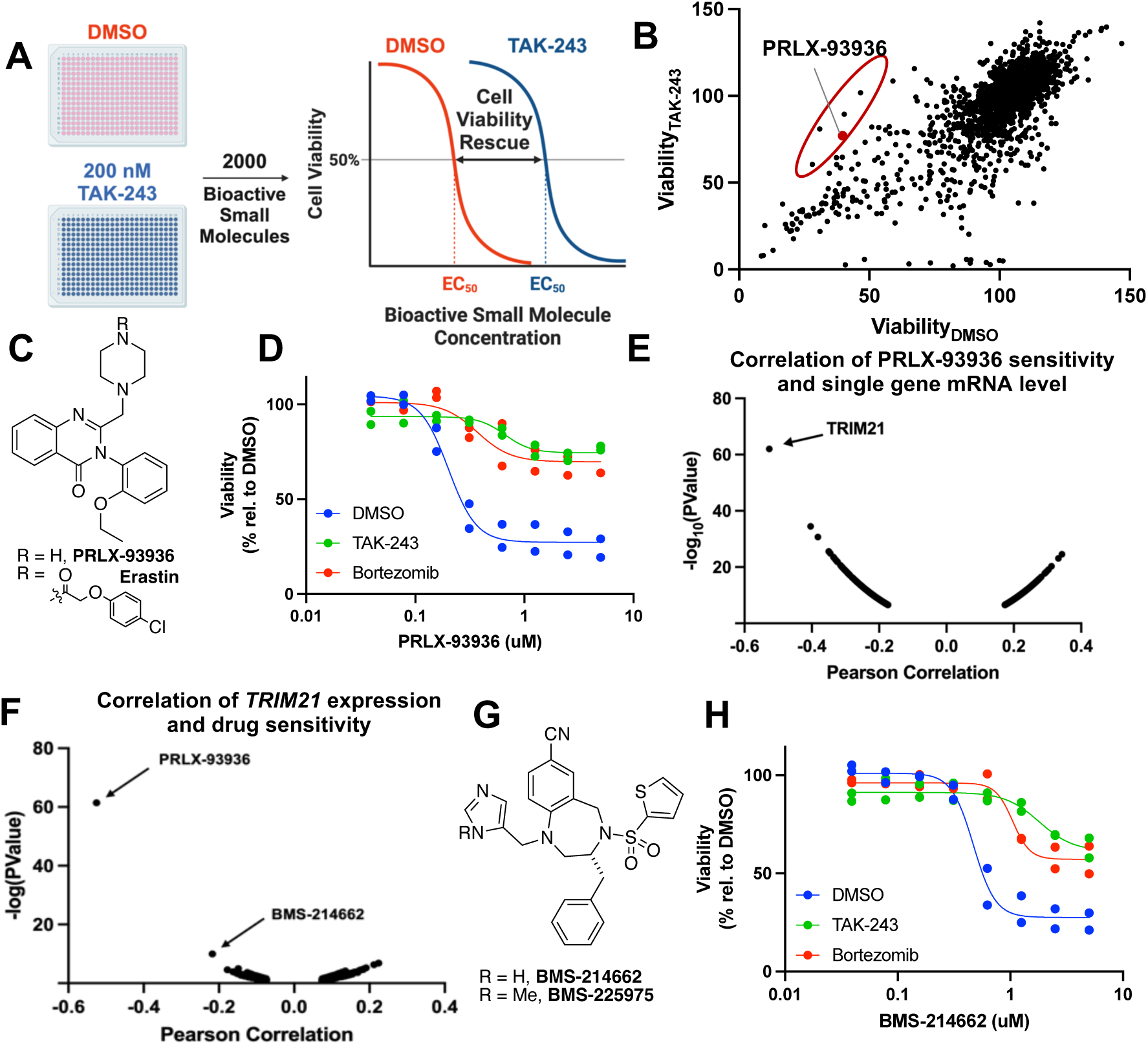
A synthetic rescue screen identifies PRLX-93936 as inducing cell death via a ubiquitin-dependent mechanism. **A)** Schematic of the synthetic rescue screening strategy. Desired molecules show reduction of cytotoxicity when ubiquitin E1 ligase activity is impaired by inhibition with the UAE inhibitor TAK-243, indicative of a mechanism of cell death reliant on ubiquitination. **B)** Dot-plot representing the performance of each 2,000 bioactive small molecules in the synthetic rescue screen performed using OCI-AML-3 cells treated for 10 h. X-axis, viability measured for each library molecule with only DMSO vehicle treatment; y-axis, viability measured for each library molecule co-treated with TAK-243 (200 nM). Hits (red oval) were significantly cytotoxic as single agents (viability < 50%, x-axis) and showed at least 30% greater viability in combination with TAK-243. Red circle, PRLX-93936. **C)** Structures of PRLX-93936 and erastin. **D)** Cell viability (CellTiter-Glo) following treatment with PRLX-93936 alone or in combination with TAK-243 (200 nM) or Bortezomib (500 nM) for 10 h in OCI-AML-3 cells (n=2 independent experiments, each with at least 2 wells per concentration). **E,F)** Correlation plots of publicly-available cancer cell line profiling data (www.depmap.org) establishing that TRIM21 expression is correlated with PRLX-93936 cytotoxicity (e) and that the cytotoxicity of PRLX-93936 and BMS-214662 is uniquely well correlated with TRIM21 among the small molecules within the database (f). **G)** Structures of BMS-214662 and BMS-225975. **H)** Cell viability (CellTiter-Glo) following treatment with PRLX-93936 alone or in combination with TAK-243 (200 nM) or Bortezomib (500 nM) for 10 h in OCI-AML-3 cells (n=2 independent experiments, each with at least 2 wells per concentration).

PRLX-93936 is a cytotoxin of unknown mechanism of action that has been linked to diminished HIF pathway signaling.^19,20^ Publicly-available cancer cell line profiling data (depmap.org) indicated that PRLX-93936 induced wide-ranging cytotoxic responses in cancer cell lines, with many potently killed and others fully resistant (Supp. Fig. 1c). Correlation of cell killing potency with expression of single transcripts has previously provided mechanistic understanding for molecules of unknown mechanism-of-action, including the molecular glue CR-8.^4,33^ Analysis of publicly-available transcriptomic profiles revealed that sensitivity to PRLX-93936 was uniquely correlated with high expression of TRIM21, a member of the TRIM family of E3 ligases (Fig. 1e, Supp. Fig. 1c). Conversely, among the thousands of small molecules in the DepMap database, PRLX-93936 was best-correlated with TRIM21 transcript levels, highlighting the strong association between PRLX-93936 and TRIM21 (Fig. 1f).

A second molecule, the farnesyl transferase inhibitor (FTI) BMS-214662, also appeared well-correlated with TRIM21 expression level (Fig. 1f,g). Interestingly, work from Bristol Myers Squibb and others has established that this molecule has an additional uncharacterized apoptotic mechanism not observed for closely related FTI inhibitors, including its *N*-methyl analog BMS-225975.^34–36^ We next confirmed that BMS-214662, like PRLX-93936, induced rapid cell death in OCI-AML-3 cells that could be suppressed by co-treatment with either TAK-243 or bortezomib (Fig. 1h). Together these data suggested that PRLX-93936 and BMS-214662 share a cytotoxic mechanism of action that involves TRIM21-mediated ubiquitination.

We next confirmed strong expression of TRIM21 in both Jurkat and OCI-AML-3 cells and used CRISPR/Cas9 targeting to knockout TRIM21 in both cell lines (Fig. 2a,b). Strikingly, while WT cells were highly sensitive to PRLX-93936 (EC_50_ ca. 100 nM), TRIM21 KO in both Jurkat and OCI-AML-3 cells led to full resistance to PRLX-93936 at concentrations as high as 50 μM (Fig. 2c,d). Likewise, both TRIM21 KO cell lines showed >100-fold increase in EC_50_ for BMS-214662 (Fig. 2e,f). Notably, while PRLX-93936 shares some structural features with erastin, an inducer of ferroptosis, erastin sensitivity was unaffected by TRIM21 KO (Fig. 2g). Likewise, the cytotoxicity of farnesyl transferase inhibitor BMS-225975, which differs from BMS-214662 by only addition of a methyl group, was also unaffected by TRIM21 KO (Fig. 2h).

**Figure 2.**
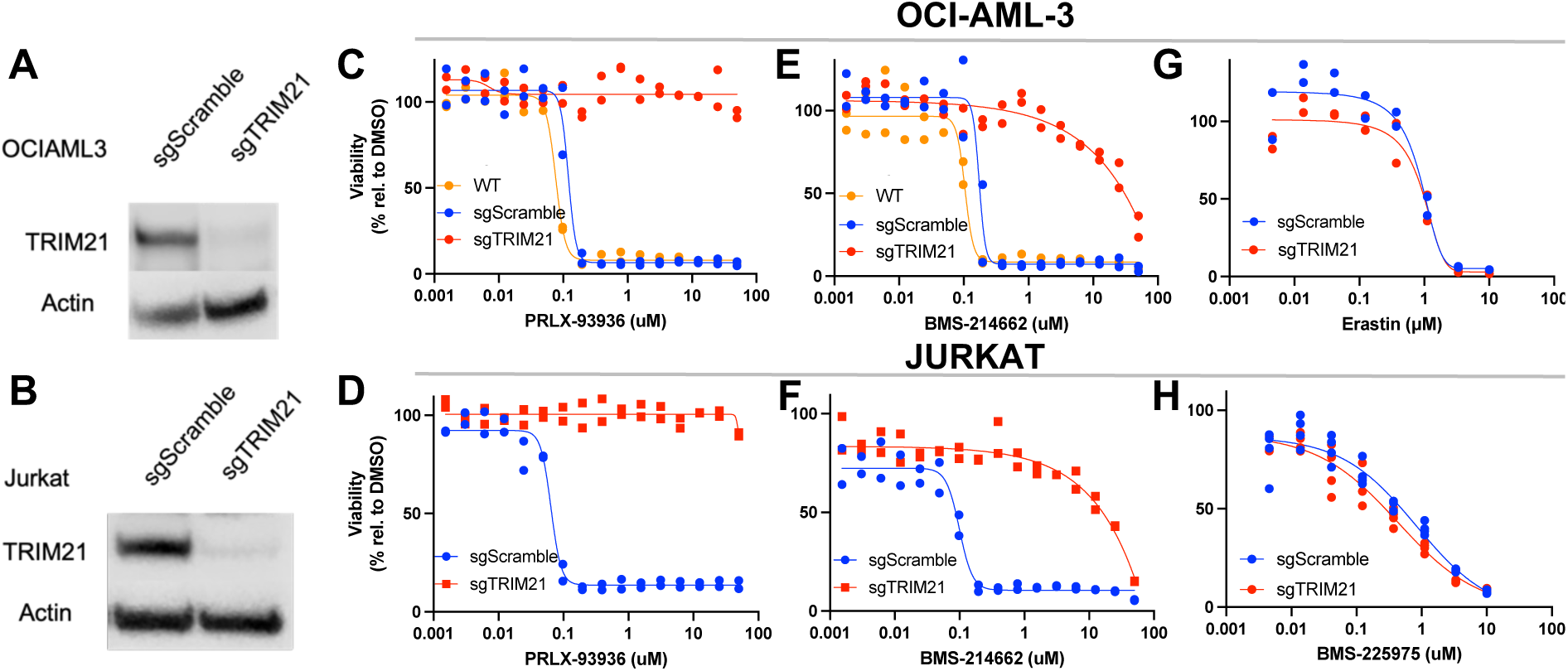
TRIM21 loss prevents cell killing by PRLX-93936 and BMS-214662. **A,B)** Western blot following CRISPR/Cas9 targeting of TRIM21 in OCI-AML-3 cells (A) and Jurkat cells (B). **C,D)** Cell viability following treatment with PRLX-93936 for 24 h in OCI-AML-3 cells (C) or Jurkat cells (D). **E,F)** Cell viability measurements as described for C,D but using BMS-214662. **G)** Cell viability following treatment with erastin for 72 h in OCI-AML-3 cells. **H)** Cell viability following treatment with BMS-225975 for 72 h in Jurkat cells. All cell viability measurements used CellTiter-Glo. All CTG data represents n=2 independent experiments, each with at least 2 wells per condition, except h which is n=1 with 2 wells per condition.

We also evaluated the impact of overexpression of TRIM21 on PRLX-93936 and BMS-214662 sensitivity. OCI-AML-3 cells expressing a TRIM21-FLAG allele became ca. 10-fold more sensitive to both PRLX-93936 and BMS-214662 (Fig. 3a-c). In contrast, expression of an established catalytically inactive triple Cys-to-Ala TRIM21 mutant (TRIM21-CA) partially suppressed the cell killing induced by these probes (Fig. 3a-c).^37,38^ Since TRIM21 enzymatic activity is known to be enhanced by its oligomerization, this catalytically inactive allele may induce a dominant-negative effect and partially mimic TRIM21 loss-of-function. We also overexpressed TRIM21 in C33A cells, a cervical cancer line that both publicly available transcriptomics data (www.depmap.org) and immunoblotting support as not expressing TRIM21 (Fig. 3d). While parental C33A cells were resistant to PRLX-93936 and BMS-214662 at concentrations up to 10 μM, overexpression of TRIM21 greatly sensitized C33A cells to these probes, with EC_50_ values comparable to those seen in WT OCI-AML-3 or Jurkat cells (Fig. 3e,f). As expected, expression of the catalytically inactive TRIM21 CA mutant allele had no impact on sensitivity (Fig. 3e,f). As a final example, overexpression of TRIM21 in HEK293T cells also induced extraordinary sensitivity to PRLX-93936 and BMS-214662 (Fig. 3g-i). Together these gain- and loss-of-function genetic manipulations indicate that TRIM21 catalytic activity is both necessary and sufficient for PRLX-93936 and BMS-214662 to induce cell death.

**Figure 3.**
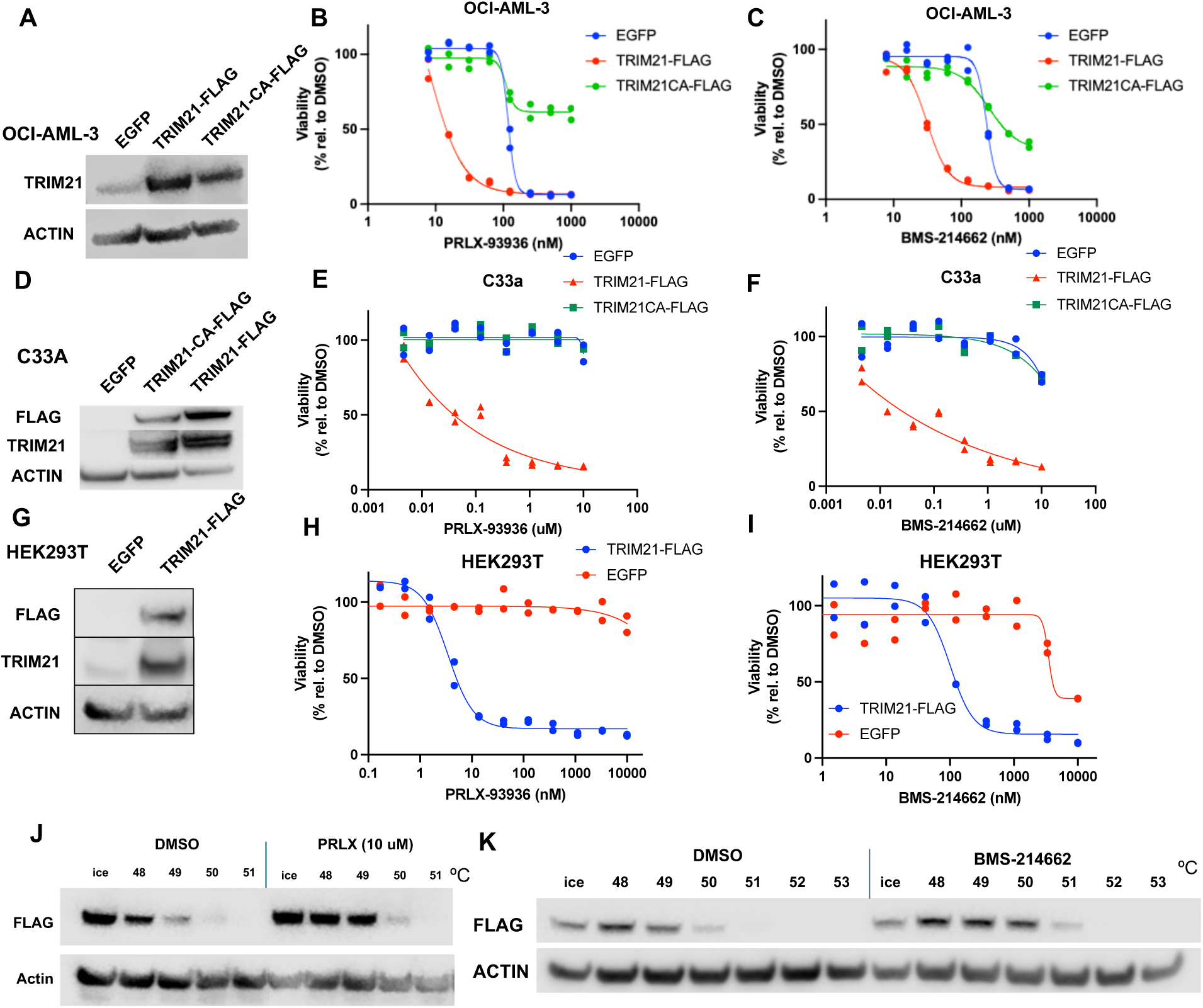
Overexpression of TRIM21 drives sensitivity to PRLX-93936 and BMS-214662. **a,d,g)** Western blot following lentiviral overexpression of TRIM21-FLAG or inactive mutant TRIM21-CA-FLAG in OCI-AML-3 cells (a), C33A cells (d), or HEK293T cells (g)(representative image from two independent experiments). **b,e,h)** Cell viability following treatment with PRLX-93936 for 24 h in TRIM21-expressing OCI-AML-3 cells (B) or 72 h in TRIM21-expressing C33A (e) or HEK293T cells (h) (n=2 independent experiments, each with at least 2 wells per concentration). **c,f,i)** Cell viability following treatment with BMS-214662 for 24 h in TRIM21-expressing OCI-AML-3 cells (C) or 72 h in TRIM21-expressing C33A (f) or HEK293T cells (i) (n=2 independent experiments, each with at least 3 wells per concentration). j,k) CETSA experiment monitoring the impact of PRLX-93936 (j; n = 1) or BMS-214662 (k; representative of two independent experiments) on the thermal denaturation of FLAG-TRIM21 in OCI-AML-3 cells.

To evaluate whether these small molecules directly bind TRIM21, we performed cellular thermal shift (CETSA).^39^ PRLX-93936 induced clear stabilization of TRIM21-FLAG by 1-2 °C, consistent with a direct binding interaction (Fig. 3j). BMS-214662 stabilized TRIM21-FLAG by a similar margin, providing additional evidence that these molecules induce cell death by direct binding to TRIM21 and subsequent ubiquitination and proteasomal degradation of an essential protein (Fig. 3k). As existing TRIM21-targeting glues have been demonstrated to bind to TRIM21’s PRYSPRY domain, we hypothesize that PRLX-93936 and BMS-214662 also likely bind within this established druggable pocket.^29,30^

To identify the protein or proteins degraded by PRLX-93936 and BMS-214662, we next performed an unbiased proteomics analysis. We initially evaluated PRLX-93936 treatment in both Jurkat and OCI-AML-3 cells to identify proteins comparably depleted across these equally sensitive cell lines. Strikingly, a cluster of nucleoporin proteins were strongly downregulated in both Jurkat and OCI-AML-3 cells (Fig. 4a,b, Supporting Fig. 2a,b). Strongest effects were seen for Nucleoporin 214 (NUP214) and Nucleoporin 88 (NUP88), which are essential proteins known to directly interact at the cytoplasmic face of the nuclear pore (Fig. 4b,d).^40,41^ However, numerous additional essential nucleoporin proteins–NUP98, NUP188, NUP155, and NUP35, among others–were also reduced to a lower extent (Fig. 4a, Supporting Fig. 2c). Importantly, changes in NUP214 and NUP88 levels were not observed when the proteomics experiment was repeated using TRIM21 KO OCI-AML-3 cells, indicating that the reduction of these nucleoporins is TRIM21-dependent (Fig. 4c-e, Supporting Fig. 2c,d). Additionally, proteomic analysis revealed that nucleoporin degradation was rapid, with levels of some nucleoporins reduced by half within an hour (Fig. 4f).

**Figure 4.**
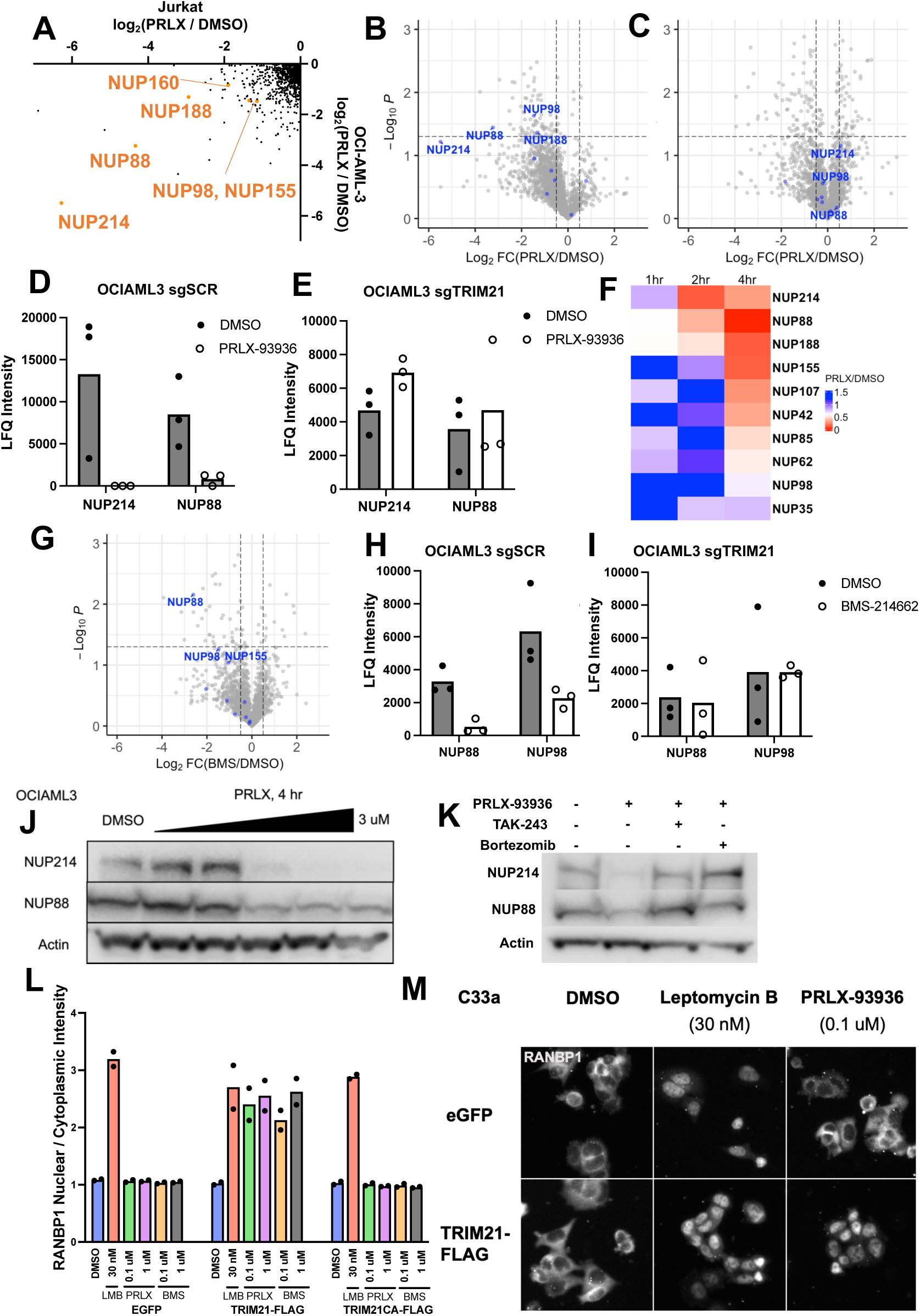
PRLX-93936 and BMS-214662 induce degradation of nucleoporin protein and impair nuclear transport. **a)** Label-free LC/MSMS quantitation of reductions of protein levels following PRLX-93936 treatment (500 nM, 6 h) in Jurkat cells (x-axis) and OCI-AML-3 cells (y-axis), with alterations in nucleoporin proteins highlighted in orange. **b,c)** Volcano plot highlighting alterations in protein levels following treatment with PRLX-93936 as in a in OCI-AML-3 sgSCRAMBLE (sgSCR) cells (b) and OCI-AML-3 sgTRIM21 cells (c) with nucleoporin proteins labeled in blue. **d,e)** LFQ intensity values for specific nucleoporins noted panels a-c in both sgSCR and sgTRIM21 OCI-AML-3 cells. **f)** Heatmap representing the fold change in nucleoporin abundance as measured by label-free LC/MSMS quantitation after treatment with PRLX-93936 (500 nM) for 1, 2, or 4 hours. **g)** Volcano plot as in b except OCI-AML-3 cells are treated with BMS-214662 (1 μM) for four hours. **h,i)** LFQ intensity values for specific nucleoporins in both sgSCR and sgTRIM21 OCI-AML-3 cells following treatment with BMS-214662 (1 μM) for four hours. **j)** Western blot demonstrating reduction of NUP214 and NUP88 protein levels with increasing concentrations of PRLX-93936 treatment (4 hours, OCI-AML-3 cells). **k)** Western blot evaluating the impact of pre-treatment (2 hours) with the proteasome inhibitor bortezomib (5 μM) or UAE inhibitor TAK-243 (0.5 μM) prior to addition of PRLX-93936 (1 μM) for 4 hours. **l)** Quantitation of immunofluorescence imaging of RANBP1 reported as the ratio of nuclear to cytoplasmic signal intensity in the indicated three C33A overexpression cell lines. ‘PRLX’, PRLX-93936; ‘BMS’, BMS-214662. All treatments 6 hours. Points represent average of 2 wells/condition with at least 900 cells quantified/condition from n=2 independent experiments. **m)** Representative images of RANBP1 subcellular localization following the treatments in l.

We then repeated these proteomics experiments with BMS-214662. As with PRLX-93936, multiple nucleoporins were degraded by BMS-214662 after 4 h in OCI-AML-3 cells, with NUP88 and NUP98 most strongly affected (Fig. 4g,h). Critically, nucleoporin levels were unaffected by BMS-214662 treatment in OCI-AML-3 cells lacking TRIM21, further supporting TRIM21’s essential role in mediating nucleoporin degradation (Fig. 4i, Supp. Fig. 2e). Moreover, BMS-225975, which differs from BMS-214662 only by addition of a methyl group but lacks TRIM21-dependent cell killing (Fig. 2h), had no impact on nucleoporin levels (Supp. Fig. 4f).

We also used western blotting as an orthogonal approach to validate our proteomics results. Levels of both NUP214 and NUP88 were substantially depleted following treatment with PRLX-93936 for 4 h in OCI-AML-3 cells (Fig. 4j). In contrast, reductions in NUP214 and NUP88 were not observed when PRLX-93936 was co-treated with E1 inhibitor TAK-243 or proteasome inhibitor bortezomib, indicating the nucleoporin loss was ubiquitin- and proteasome-dependent (Fig. 4k).

Loss of numerous essential nucleoporin proteins would be expected to impair nuclear trafficking. We next evaluated nuclear export in C33A cells using immunofluorescence detection of RANBP1 as a common marker of nuclear-to-cytoplasmic trafficking. C33A cells express TRIM21 at undetectable levels and are resistant to PRLX-93936/BMS-214662 unless TRIM21 is overexpressed (Fig. 3). As expected, in C33A cells, PRLX-93936 and BMS-214662 had no impact on the subcellular localization of RANBP1, while a canonical nuclear export inhibitor, the XPO1-targeting natural product Leptomycin B, led to strong nuclear accumulation of RANBP1 (Fig. 4l,m). In contrast, treatment of TRIM21-overexpressing C33A cells with multiple concentrations of either PRLX-93936 or BMS-214662 induced nuclear accumulation of RANBP1 comparable to Leptomycin B treatment (Fig. 4l,m). However, nuclear accumulation of RANBP1 was not observed in cells overexpressing catalytically inactive TRIM21 (Fig 4l). Importantly, this TRIM21-dependent nucleoporin degradation mechanism helps to rationalize past unexplained observations of extensively altered subcellular protein localization following BMS-214662 treatment.^36^ Together, these data highlight that the observed loss of multiple essential nucleoporins leads to lethal nuclear trafficking deficits that are dependent on catalytically active TRIM21.

In the course of this work, hydroxy-acepromazine was reported as a small molecule that induces cancer cell death via TRIM21-mediated degradation of nucleoporins (Fig. 5a).^29^ These studies also provided genetic evidence that hydroxy-acepromazine induced degradation of multiple nucleoporins by recruiting TRIM21 specifically to NUP98. We confirmed that hydroxy-acepromazine indeed caused TRIM21-dependent cell death in OCI-AML-3 cells, albeit with 100-fold diminished potency relative to PRLX-93936 and BMS-214662 (Fig. 5b,c). Additionally, we confirmed that targeted disruption of the autoproteolysis domain of NUP98 was sufficient to prevent cell killing by PRLX-93936 and BMS-214662. Delivery of recombinant Cas9 and five independent single guide RNAs (sgRNAs) targeting this region of NUP98 led to isolation of A549 cells that were strongly resistant to both PRXL-93936 and BMS-214662 (Fig. 5d-e). These resistant cells did not show resistance to the unrelated cytotoxic agent TAK-243, supporting a selective effect (Fig. 5f). Each of these five distinct sgRNAs led to detection of a 200 kDa NUP98 proteoform in these PRLX-93936-resistant cells, indicating that the autoproteolysis of NUP98 had been impaired (Fig. 5g). Impaired autoproteolysis of NUP98 was recently demonstrated as a resistance mechanism to hydroxy-acepromazine.^29^ Finally, these PRLX-93936-resistant cells did not show reduction in NUP214 or NUP88 levels following treatment with PRLX-93936, providing additional support for nucleoporin degradation as essential for this cytotoxic mechanism (Fig. 5h).

**Figure 5.**
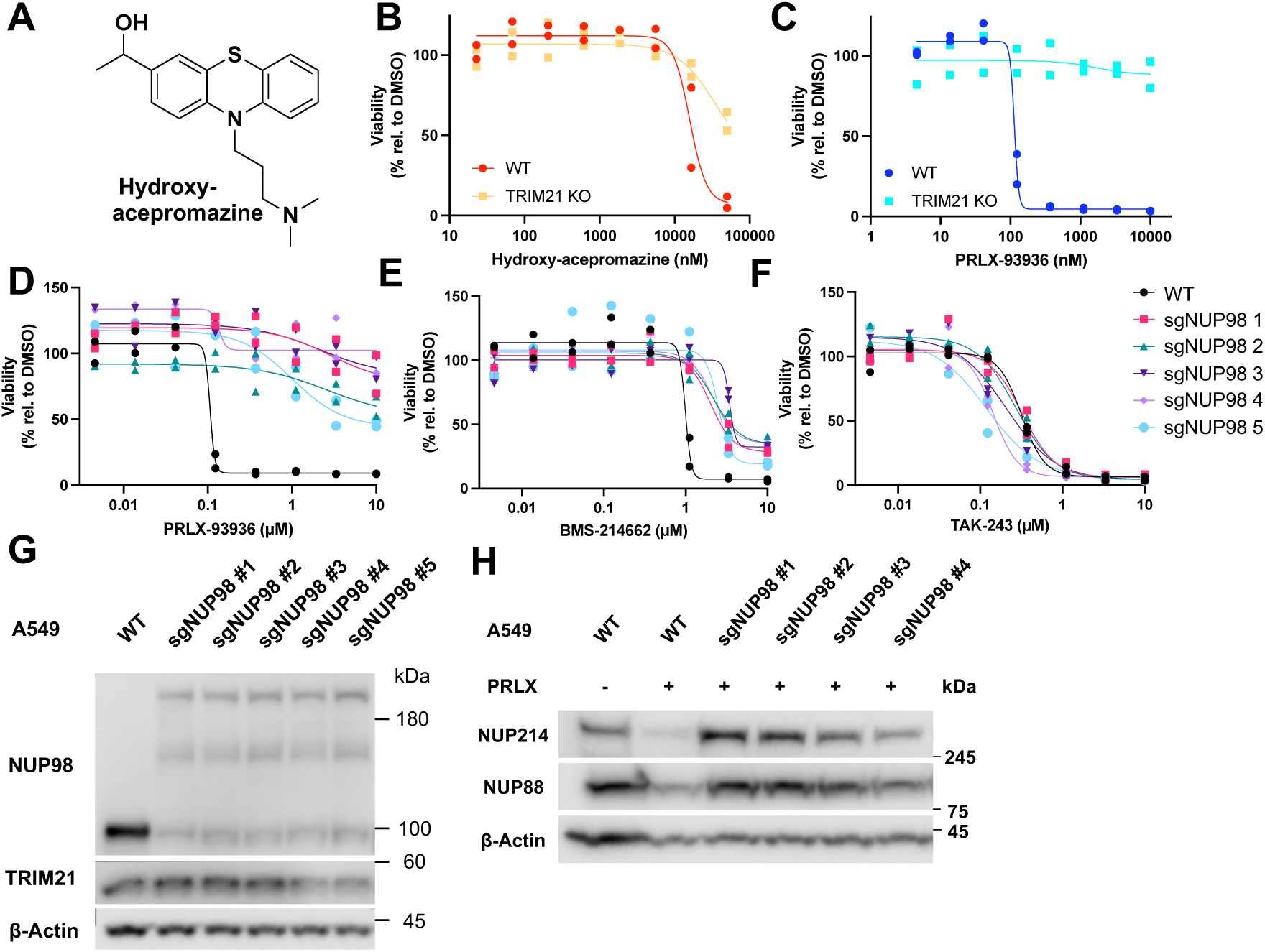
Like hydroxy-acepromazine, PRLX-93936 and BMS-214662 phenotypes can be abrogated by targeting the NUP98 autoproteolysis domain. a) Structure of hydroxy-acepromazine. b,c) Viability (CellTiter-Glo) of OCI-AML-3 cells (WT or TRIM21 KO) following 24 h treatment with the indicated concentrations of hydroxyacepromazine (b) or PRLX-93936 (c). d-f) A549 cells were subjected to CRISPR/Cas9 targeting using 5 sgRNAs targeting the autoproteolysis domain of NUP98 and then exposed to PRLX-93936 (1 μM) for 7 days. Surviving cells were then exposed to the indicated concentrations of PRLX-93936 (d), BMS-214662 (e) or TAK-243 (f) and cell viability was measured after 72 h with CellTiter-Glo. All CellTiter-Glo experiments represent n=2 independent experiments with at least 2 wells per condition. g) Western blot of NUP98 in WT A549 as well as A549 cells targeted with the five independent NUP98 sgRNAs. A NUP98 proteoform indicative of impaired autoproteolysis is highlighted. h) Western blot of NUP214 and NUP88 following PRLX-93936 treatment (1 μM /12 hr) in A549 cells resistant to PRLX-93936 following treatment with four of the NUP98 sgRNAs reported in d-g.

## Discussion

We have established that PRLX-93936 and BMS-214662 induce cell death via TRIM21-mediated degradation of nucleoporins. Genetic manipulations support TRIM21 as necessary and sufficient for the observed cell death across multiple cell lines. Extensive proteomic analyses identified a wide range of nucleoporins as degraded following treatment with PRLX-93936 and BMS-214662, and multiple sgRNAs targeting NUP98’s autoproteolysis domain provided strong resistance to these small molecules. This finding establishes that NUP98 plays an essential role in the observed nucleoporin degradation and cell death and may directly contact TRIM21 in the presence of PRLX-93936 and BMS-214662. Notably, NUP98 levels were unchanged in TRIM21 KO cells (Supporting Fig. 2), suggesting that TRIM21 does not regulate NUP98 levels in the absence of small molecule treatments.

Very recent work has established other small molecules that induce cell death via TRIM21-mediated nucleoporin degradation, including hydroxy-acepromazine^29^ and HGC652.^30^ These molecules, like PRLX-93936 and BMS-214662, share the ability to degrade nucleoporins as well as many other proteins, highlighting how loss of nuclear pore function can induce broad and rapidly lethal disruption of the proteome. Notably, GLE1, an mRNA export factor strongly depleted by hydroxy-acepromazine and HGC652degraders, was also depleted to undetectable levels in PRLX-treated OCIAML3 cells, further supporting a convergent mechanism for these molecules. Interestingly, these four series of TRIM21-targeting glues share almost no structural similarity, highlighting how distinct scaffolds can equally function to induce nucleoporin degradation. Notably, PRLX-93936 and BMS-214662 offer substantially enhanced potency relative to hydroxy-acepromazine (Fig. 5) and appear comparable to HGC652.

A unique aspect of PRLX-93936 and BMS-214662 is that these molecules have previously been investigated clinically in cancer. While clinical data for PRLX-93936 have only been presented at conferences,^42,43^ published reports indicate that BMS-214662 was well tolerated and showed robust objective responses in a subset of acute myeloid leukemia and myelodysplastic syndrome patients.^44^ Apoptotic responses were observed in tumors following treatment with BMS-214662 but not other farnesyl transferase inhibitors, suggesting that the observed clinical responses were unlikely a result of farnesyl transferase inhibition. Together with the observation that high expression of TRIM21 strongly predicts PRLX-93936 and BMS-214662 sensitivity (Fig. 1), these findings suggest the potential for clinical re-evaluation of these agents in subsets of patients whose tumors have high TRIM21 expression.

Previous work with hydroxy-acepromazine has demonstrated that TRIM21-targeting glues can be evolved to PROTACs that degrade proteins engineered to reside within nuclear condensates.^29^ Separate from the potential of PRLX-93936 and BMS-214662 as anticancer agents, these scaffolds may also serve as starting points for elaboration to aggregate-targeting PROTACs. The design strategies and scope of aggregation-prone proteins amenable to TRIM21-mediated degradation remain to be established; however, TRIM21’s unique biological preference for ubiquitination of antibody-coated pathogens and other large aggregates suggests that TRIM21 may be ideally suited for degrading aggregated proteins. Establishing PRLX-93936 and BMS-214662 as highly potent TRIM21-targeting molecular glues opens new opportunities for leveraging TRIM21 both for cancer therapy and toward a wider range of PROTACs and molecular glues.

## Acknowledgments

This work was supported by support from the CWRU School of Medicine and Thomas F. Peterson, Jr. M.A.S and P.Z. were supported by the CWRU Medical Scientist Training Program (NIH T32GM007250). Additional support was provided by the Small-Molecule Drug Development and Proteomics Shared Resources of the Case Comprehensive Cancer Center (P30 CA043703). The timsTof Pro2 instrument used for LC-MS/MS was purchased via an NIH shared instrument grant, S10 OD030398. We thank B. Willard, L. Li, Y.F. Chen, D. Taylor, W. Huang, E. Schick, and F. Najm for technical assistance and discussion.

## Author Contributions

M.A.S., Y.F., and D.J.A. performed screening and analysis of bioactive small molecules. M.A.S. led experimental efforts to characterize PRLX-93936 and BMS-214662. E.H., A.R.A., P.Z., R.B.G., and A.N.F. performed cell-based assays. M.A.S. and D.J.A. wrote the manuscript. All authors participated in data analysis and manuscript editing and approved the final manuscript.

## Declaration of Interests

D.J.A. and M.A.S. are inventors on a provisional patent application filed by Case Western Reserve University related to this work.

## SUPPORTING FIGURES

**Figure S1.**
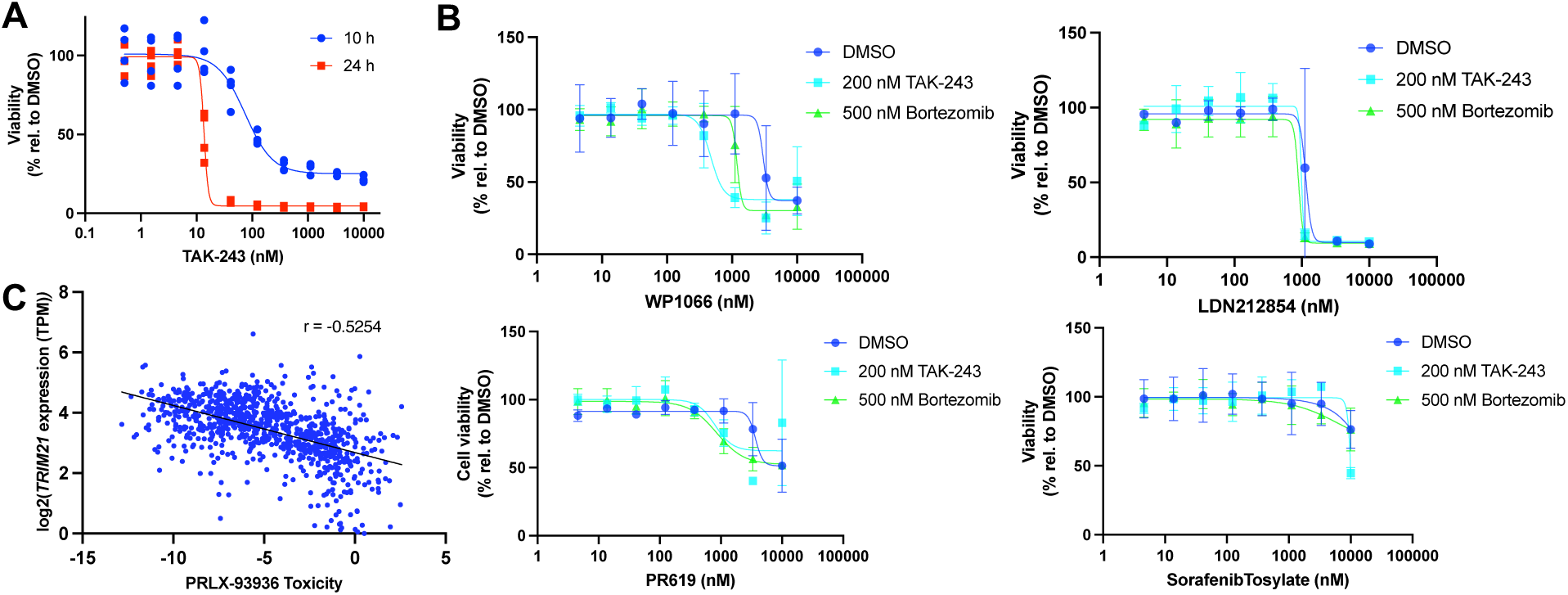
**A)** Cell viability (CellTiter-Glo) following treatment of OCI-AML-3 cells with TAK-243 at the indicated concentrations for either 10 h (blue) or 24 h (red). These data informed selection of 20 nM TAK-243 for our rescue screen in Fig. 1b. n = 1 independent experiment with 2 wells/condition **B)** Cell viability (CellTiter-Glo) for the four additional hits that proved to be false positives and did not show suppression of cytotoxicity upon retest. n=2 independent experiment with 2 wells/condition **C)** Correlation of PRLX-93936 cytotoxicity (PRISM)and TRIM21 transcript level across the cancer cell lines available within DepMap (www.depmap.org). A cytotoxicity value of 0 indicates no effect, while more negative numbers reflect more potent cytotoxic effects.

**Figure S2.**
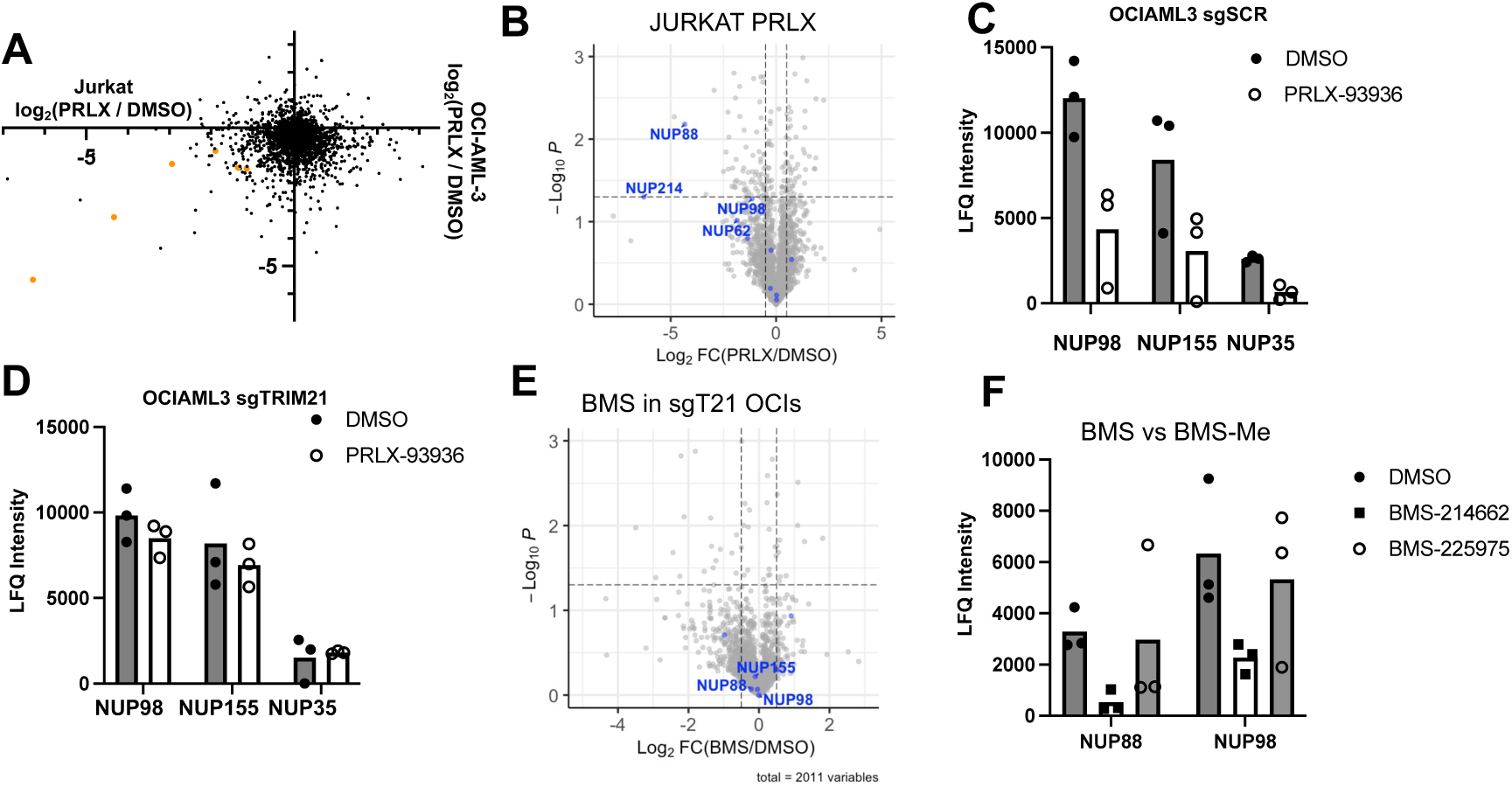
Further proteomic characterization of PRLX-93936 and BMS-214662. **a)** Label-free LC/MSMS quantitation of changes of protein levels following PRLX-93936 treatment (500 nM, 6 h) in Jurkat cells (x-axis) and OCI-AML-3 cells (y-axis), with alterations in nucleoporin proteins highlighted. This expanded version of Figure 4a shows all 4 quadrants. b) Volcano plot highlighting alterations in protein levels following treatment with PRLX-93936 (500nM, 6 hr) in Jurkat cells. c,d) LFQ intensity values for specific nucleoporins in both sgSCR (c) and sgTRIM21 (d) OCI-AML-3 cells. e) Volcano plot highlighting alterations in protein levels following treatment with BMS-214662 (1 μM, 4 h) in OCI-AML-3 TRIM21 KO cells. f) LFQ intensity values for specific nucleoporins in both OCI-AML-3 cells treated with BMS-214662 (black squares) or BMS-225975 (1 μM, 4 h), the N-methylated analog of BMS-214662 that lacks TRIM21-dependent cell killing (open circles). Each proteomics experiment represents n=1 independent experiment containing 3 technical replicates.

## METHODS

### Cell lines

Cell lines were incubated at 37 °C with 5% CO 2 under humidified conditions. Lenti-X 293T cells were purchased from Takara Bio. JURKAT E6-1 (TIB-152) and A549 (CRM-CCL-185) were purchased from ATCC. OCI-AML-3 were a gift from Dr. David Wald (CWRU). C33a were a gift from Dr. John Pink (CWRU). HEK 293T cells were a gift from Dr. Derek Taylor (CWRU). OCI-AML-3, JURKAT cell lines were maintained in glutamine-containing RPMI-1640 supplemented with 10% fetal bovine serum (FBS), and 1% penicillin/streptomycin. Lenti-X 293T, HEK 293T, and A549 cells were maintained in DMEM medium supplemented with 10% FBS,2 mM L-glutamine, and 1% penicillin/streptomycin. C33a cells were maintained in glutamine-containing EMEM medium supplemented with 10% FBS and 1% penicillin/streptomycin.

### High-Throughput Chemical Screen

OCI-AML-3 cells were seeded in a sterile tissue culture 384-well plate (Corning, 3765) at 1000 cells and 50 μl of media per well using an EL406 Microplate Washer Dispenser (Biotek). Screening library molecules (a collection of 2000 molecules derived from the Sigma LOPAC1280 and Selleck Bioactive Compound Library L1700 collections) were maintained as a 3 mM DMSO stock in Abgene storage 384-well plates (ThermoFisher Scientific; AB1055). Transfer of 100 nL of each library molecule to assay plates was achieved using a solid pin tool, resulting in a final screening concentration of 6 μM. DMSO was used as a negative control within each plate. From each library plate, two replicate assay plates were generated that were identical except one plate was co-treated with 200 nM TAK-243 (added directly to cells and media 1 hour before dispensing of library compounds). Plates were then incubated at 37°C for 10 h, at which time 5 μL of CellTiter-Glo reagent (Promega, G7572) was added to each well using the EL406 Microplate Washer Dispenser (Biotek). Plates were allowed to equilibrate for 10 min with shaking at room temperature before luminescence was immediately read using a Synergy *Neo*2Multimode microplate reader (Biotek). Viability was calculated relative to DMSO wells within each plate, and hits were called on the basis of at least 50% reduction in viability in the DMSO conditions as well as a % viability gain of at least 30% in the TAK-243 condition.

### Cell Viability Assay

OCI-AML-3 and JURKAT cells were plated at a density of 1000 cells in 50 μl of culture medium in 384-well plates and treated with inhibitors same day before incubation at 37°C. For C33a, A549, and HEK 293T cells, 500 cells in 50 μl in 384-well plates were used. After 24 h (OCI-AML3, JURKAT) or 72 h (C33a, A549, HEK 293T), 5 μl of CellTiter-Glo (Promega) was added to each well, and the contents were mixed on an orbital shaker at 25 °C for 15 min. Luminescence was recorded using a Biotek Synergy Neo2 microplate reader with Gen5 3.03.14 software.

### Lentivirus Production and Stable Cell Line Generation

Lentiviral ORF-containing plasmids were synthesized by Vectorbuilder. Lenti-X 293T cells were plated in 6 well plates and allowed to attach overnight at 37°C. Cells were then co-transfected with lentiviral plasmids, psPAX2, and pMD2.G using Lipofectamine 2000 (Invitrogen, 11668027) per manufacturer’s instructions. After 6 hours, media was replaced, and cells were incubated for 48hr. Lentivirus containing media was collected and filtered through a 0.45 μm filter before being directly added to cells with 8 ug/mL polybrene (Sigma Aldrich TR-1003-G). After 24 hrs at 37°C, media was replaced, and cells were incubated for another 24 hrs before selection with puromycin or blasticidin.

### Western Blotting

Cells were washed twice with PBS and lysed in Pierce RIPA Buffer (Thermo Fisher Scientific, 89900) supplemented with 1X cOmplete^™^ Mini EDTA-free Protease Inhibitor Cocktail (Roche, 11836170001). OCI-AML-3 cells were washed twice with PBS and resuspended in PBS supplemented with 1X cOmplete^™^ Mini EDTA-free Protease Inhibitor Cocktail before being lysed with two freeze-thaw cycles in liquid nitrogen. Lysates were clarified by centrifugation at 20,000xg for 20 min at 4°C. Soluble protein concentrations were quantified with a Pierce BCA Assay (Thermo Fisher Scientific, 23225). Lysates were normalized by protein concentration and separated on 4-12% or 3-8% gradient gels (Invitrogen, NW04125BOX or TA03815BOX) before being transferred to PVDF membranes using the XCell II Blot Module (Thermo Fisher Scientific, EI9051). Membranes were blocked in 5% milk in TBST and incubated in primary antibody overnight at 4°C. Primary antibodies used: anti-β-actin (Sigma-Aldrich, A3854, 1:10,000), anti-TRIM21 (Proteintech, 12108-1-AP, 1:1000), anti-FLAG M2 (Sigma-Aldrich, F3165), anti-NUP214 (Abcam, ab70497, 1:500), anti-NUP88 (Santa Cruz Biotechnology, sc-365868), anti-NUP98 (Abclonal, A0530). Membranes were then incubated with secondary antibody conjugated to horseradish peroxidase (Cell Signaling Technology, 7076 or 7074, 1:1000). Membranes were developed using either SuperSignal West Pico Plus Chemiluminescent Substrate (Thermo Fisher Scientific, 34580) or SuperSignal West Femto Maximum Sensitivity Substrate (Thermo Fisher Scientific, 34095) before imaging using a LI-COR Odyssey Fc Imaging System.

### Liquid chromatography–tandem mass spectrometry

After incubation, cells were washed twice with PBS and resuspended in PBS supplemented with 1X cOmplete^™^ Mini EDTA-free Protease Inhibitor Cocktail (Roche, 11836170001). Cells were lysed with a Fisher Scientific Sonic Dismembrator Model 60 using 15 × 1-s pulses at a power level of 3 at 4 °C. Protein concentrations were quantified using a Pierce BCA Protein Assay kit (Thermo Fisher Scientific, 23225), and colorimetric development was measured using a Biotek Synergy Neo2 microplate reader. 50 ug of protein was denatured in 8 M Urea, reduced with 10 mM dithiothreitol, and alkylated with 25 mM iodoacetamide. Samples were then diluted with 100 mM Ammonium Bicarbonate and digested with 1 ug of Lys-C (Promega V1671) overnight at room temperature. Peptides were desalted on C18 reverse-phase spin columns (BioPureSPN Mini FastEq, TARGA C18, 120Å, The Nest Group, HUM S18R) before elution with 0.1% formic acid in 80% acetonitrile and 20% water. Peptides were dried and reconstituted in 0.1% formic acid in water for LC-MS/MS analysis.

Samples were analyzed by LC-MS using a timsTof Pro2 instrument (Bruker) equipped with a NanoElute UHPLC system. A 5 µl aliquot of each digest was injected onto a ThermoScientific (0.5 x 5mm) Acclaim Pepmap C_18_, 5-μm, trapping column. Liquid chromatographic elution was performed using a flow rate of 0.3 μl/min on a reverse phase column (ReproSil AQ C18, 75 μm x 150 mm, 1.9-μm 120-Å). Peptide elution was performed using a binary gradient of mobile phase A, 0.1% formic acid and mobile phase B, 0.1% formic acid in acetonitrile. Each sample was analyzed using a linear gradient stating at 2% at 0 minutes and increasing to 35%B in 50 minutes, followed by an increase to 90% B in 2 minutes, then holding at 90% for 5 minutes before re-equilibration at 2% B. The electrospray voltage was 1500V. A PASEF-DDA method was utilized for peptide identification. MS1 scans were carried out with a resolution of 30,000 measuring masses between 100-1700 Da with 1/k0 values between 0.6 and 1.6vs/cm2. MS2 scans between 100 and 1700 Da at a resolution of 30,000 was performed on precursor between charge states of 2-5 with targeted intensities of 20000, an intensity threshold of 2500 au and 10 PASEF MS/MS scans were performed in each cycle with cycle times of 1.2 sec. The peptides were fragmented using CID with an isolation window of 2 Da and collision energies ranging from 20 eV (1/k0 value of 0.6 Vs/cm2) to 59 eV (1/k0 value of 1.6 Vs/cm2). Dynamic exclusion was enabled for 30 sec.

The LC-MS/MS data were searched against the human SwissProtKB database downloaded on 3-23-2022 (26,576 entries) using the program PEAKSOnline v11. These searches were performed considering full LysC peptides with no more than 2 missed cleavage sites. The MS1 and MS2 mass accuracies was set to 20ppm and 0.05 Da respectively, carbamidomethylating was considered as a fixed modification, and oxidation of methionine and protein acetylation were considered as variable modifications. Deep learning boost was enabled. The results were filtered using a reverse decoy database strategy using percolator and the PSM, Peptide, and protein FDR rates were set to 1%. Positively identified proteins were required to be identified by a minimum of 2 peptides with at least one of these being a unique peptide identification.

Label free quantitation was performed PeaksOnline using ID-directed LFQ. Match between runs was enabled with a mass tolerance of 20 ppm, retention time shift set to auto, and a 1/k0 tolerance of 0.05. Quantitation was performed on unique peptides with a feature intensity of at least 150. The LFQ output was uploaded into Perseus V2.0.11 and the intensity values were log2 transformed and the data was filtered to remove proteins with <2 quantitative values in at least one of the groups. After matrix reduction, missing values were imputed using a normal distribution with parameters of width = 0.3, downshift = 1.8, and mode = separately for each column. The LFQ intensities were used to determine the abundance ratios of the protein across groups and significance was determined using a t-test derived p-value.

The relative abundance of the positively identified peptides was determined using the extracted ion intensities (Minora Feature Detection node) with Retention time alignment. Both the raw intensities and the intensities normalized to total peptide intensity were extracted from PD 2.5 for relative abundance analysis. Perseus was used for matrix reduction and imputation. Only peptides identified in at least 2 of the 3 biological replicates in one of the groups were considered for quantitation and the peptide intensities were normalized to total peptide amount. Missing values were imputed using a normal distribution.

### Immunofluorescence

C33a eGFP and TRIM21-FLAG cells were plated at a density of 6,000 cells in 50 μl of medium in a black, clear-bottom 96-well plate (PerkinElmer, 50-209-9831) and allowed to attach overnight at 37°C. Cells were then treated with respective inhibitors for 6 h. Cells were fixed in 4% PFA for 20 min, washed with PBS, and stained with anti-RANBP1 (Abcam, ab97659) at a 1:500 dilution and DAPI (Sigma-Aldrich, D8417) at a 1:20,000 dilution. Cells were imaged using an Operetta High Content Imaging and Analysis System (Perkin Elmer), with eight fields captured per well and two wells per condition at ×20 magnification. Images were analyzed using Harmony software on the Columbus data server (PerkinElmer). In brief, live cells were identified and their nuclear regions established using DAPI staining. Around these nuclear regions, the cytoplasmic region was defined according to the region of RANBP1 staining not overlapping with the nucleus. The nuclear-to-cytoplasmic ratio of total signal intensity per well for RANBP1 was calculated based on these criteria. Data for each molecule was normalized to DMSO wells (0%) and Leptomycin B wells (100%).

### Generating PRLX-93936-resistant cells with CRISPR–Cas9 targeting of *NUP98-96*

Short guide RNA oligonucleotides targeting exon 17 of *NUP98-96* were synthesized by Synthego with default modifications containing the following sequences (all 5’-3’): GAUUUCACUAUUGGUCGGAA, UCUCUGAUUUCACUAUUGGU, AUUGUCUCUGAUUUCACUAU, AAUGCACUCUCCUUUUUCAU, GCUAAAAUUACCAAUGAAAA, UUUCAUUGGUAAUUUUAGCA, UUAGCAAGGUCAUCCAUAGA, UUACUAUACUAUUCCAUCUA, GGUAUUAUUCUCACUAAGGU, and CACAGGUAUUAUUCUCACUA. Recombinant SpCas9-2NLS purchased from Synthego was mixed with each guide for 15 min at room temperature to allow for ribonucleoprotein complex formation (20 pmol Cas9:60 pmol single guide RNA per 1 million cells). Each mixture was added to 1 million A549 cells and delivered via electroporation using the Lonza 4D-Nucleofector system and an SF Cell Line 4D-Nucleofector kit according to the manufacturer’s instructions. After electroporation, the cells were allowed to proliferate for 72 h before selection for resistant clones using 1 μM PRLX-93936. After 7 days of selection, the surviving cells were cultured further in 1 μM PRLX-93936 and harvested for analysis by western blotting and cell viability assays.

